# An online headphone screening test based on dichotic pitch

**DOI:** 10.1101/2020.07.21.214395

**Authors:** Alice E. Milne, Roberta Bianco, Katarina C. Poole, Sijia Zhao, Andrew J. Oxenham, Alexander J. Billig, Maria Chait

## Abstract

Online experimental platforms can be used as an alternative, or complement, to lab-based research. However, when conducting auditory experiments via online methods, the researcher has limited control over the participants’ listening environment. We offer a new method to probe one aspect of that environment, headphone use. Headphones not only provide better control of sound presentation but can also “shield” the listener from background noise. Here we present a rapid (< 3 minute) headphone screening test based on Huggins Pitch (HP), a perceptual phenomenon that can only be detected when stimuli are presented dichotically. We validate this test using a cohort of “Trusted” online participants who completed the test using both headphones and loudspeakers. The same participants were also used to test an existing headphone test (AP test; Woods et al., 2017, *Attention Perception Psychophysics*). We demonstrate that compared to the AP test, the HP test has a higher selectivity for headphone users, rendering it as a compelling alternative to existing methods. Overall, the new HP test correctly detects 80% of headphone users and has a false positive rate of 20%. Moreover, we demonstrate that combining the HP test with an additional test - either the AP test or an alternative based on a beat test (BT) - can lower the false positive rate to ∼7%. This should be useful in situations where headphone use is particularly critical (e.g. dichotic or spatial manipulations). Code for implementing the new tests is publicly available in JavaScript and through Gorilla (gorilla.sc).

## Introduction

Online experimental platforms are increasingly used as an alternative, or complement, to in-lab work (Assaneo et al., 2019; Kell et al., 2018; Lavan, Knight, & McGettigan, 2019; Lavan, Knight, Hazan, et al., 2019; McPherson & McDermott, 2018; Slote & Strand, 2016; Woods & McDermott, 2018; Zhao et al., 2019). This process has been hastened in recent months by the COVID-19 pandemic. A key challenge for those using online methods is maintaining data quality despite variability in participants’ equipment and environment. Recent studies have demonstrated that with appropriate motivation, exclusion criteria, and careful design, online experiments can not only produce high quality data in a short time, but also provide access to a more diverse subject pool than commonly used in lab-based investigations (Clifford & Jerit, 2014; Rodd, 2019; A. T. Woods et al., 2015).

For auditory experiments specifically, a major challenge involves the loss of control over the audio delivery equipment and the acoustic listening environment. However, certain information can be gleaned through specially designed screening tests. Here we focus on procedures for determining whether participants are wearing headphones (including in-ear and over-the-ear versions) or listening via loudspeakers. In many auditory experiments the use of headphones is preferred because they offer better control of sound presentation and provide some attenuation of other sounds in their environment. Woods et al. (2017) developed a now widely used test to determine whether listeners were indeed using headphones, based on dichotic presentation, under the premise that participants listening over headphones, but not those listening over loudspeakers, will be able to correctly detect an acoustic target in an intensity-discrimination task (Figure 1a). In each trial the listener is presented with three consecutive 200-Hz sinusoidal tones and must determine which was perceptually the softest. Two of the tones are presented diotically: 1) the “standard” and 2) the “target” which is presented at −6 dB relative to the standard. The third tone (a “foil”) has the same amplitude as the standard but is presented dichotically, such that the left and right signals have opposite polarity (antiphase, 180°). Woods and colleagues reasoned that over headphones, the standard and foil should have the same loudness, making the target clearly distinguishable as softer. In contrast, over loudspeakers the left and right signals may interact destructively before reaching the listener’s ears, resulting in a weaker acoustic signal at both ears, and thus a lower loudness for the foil than at least the standard, and possibly also the target, causing participants to respond incorrectly.

**Figure 1.**
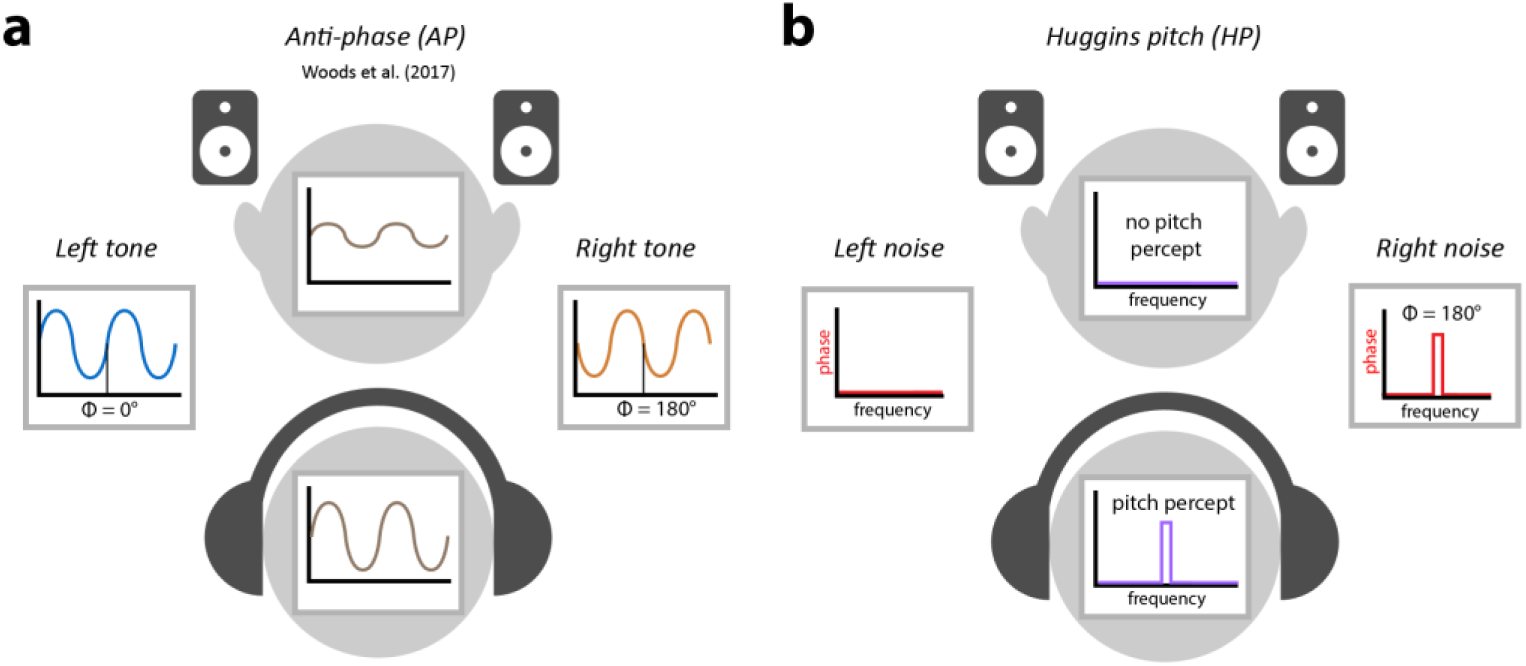
Schematic of stimuli for each test. (a) In the Woods et al test (AP test) a foil is created by presenting a 200Hz tone dichotically in anti-phase. When presented over loudspeakers this is expected to result in destructive acoustic interference, and thus reduced loudness, causing the foil to be mistaken for the target and the listener to fail the test. Over headphones there is no such interference, and thus no reduction of loudness and the listener should correctly detect the target, passing the test. However, the test is susceptible to certain loudspeaker configurations and the presence of binaural interaction which may reduce the effectiveness of the test (see text). (b) In the HP test, broadband noise is presented to one channel and the same broadband noise with a phase shift (anti-phase) over a narrow band (±6%) around 600Hz is presented to the other channel. Over headphones this results in a percept of pitch at this frequency that the listener will detect, allowing them to pass the screening. The percept depends on the left and right channels being independent, and thus tends to disappear over loudspeakers, preventing the listener from detecting the target and thus causing them to fail the test.

Woods et al. (2017) validated their test both in the lab and online. The test contained 6 trials and the threshold for passing the headphone screen was set at 5/6 correct responses. Using this threshold, in the lab 100% of participants wearing headphones passed the test, while only 18% passed when using loudspeakers. In a large number of subjects recruited online (via Amazon Mechanical Turk), only 65% passed the test, suggesting that a third of the online listeners may have actually used loudspeakers, despite instructions to use headphones. The ability to detect those cases makes this test a valuable resource.

However, this test has important limitations. Most critically, it is not strictly a test of headphone use because it is passable when listening over a single channel, e.g. if the participant is using a single earbud. Instead, the Woods et al test focuses on “weeding out” loudspeaker users by identifying the participants who are susceptible to the destructive interaction between L and R channels. This effect depends on the specific positions of the loudspeakers relative to the listener, and on other features of the space (e.g. occluders) and may not generalize to all listening environments. In particular, participants may be able to pass the test even when listening in free-field, if they are positioned in close proximity to one loudspeaker. Furthermore, the antiphase foil stimulus causes inter-aural interactions that give rise to a particular binaural percept that is not present for the other two tones, and which may be more salient over headphones. To solve the loudness discrimination task, participants must thus ignore the binaural percept and focus on the loudness dimension. This introduces an element of confusion, which might reduce performance among true headphone users. Here we present and validate a different method for headphone screening that addresses these problems.

We examine the efficacy of a headphone screening test based on a particular dichotic percept – Huggins Pitch – that should be audible via headphones but absent when listening over loudspeakers. The Huggins Pitch (HP) stimulus (Akeroyd et al., 2001; Chait et al., 2006; Cramer & Huggins, 1958; Figure 1b) is an illusory pitch phenomenon generated by presenting a white noise stimulus to one ear, and the same white noise—but with a phase shift of 180° over a narrow frequency band—to the other ear. This results in the perception of a faint tonal object (corresponding in pitch to the centre frequency of the phase-shifted band), embedded in noise. Importantly, the input to either ear *alone* lacks any spectral or temporal cues to pitch. The percept is only present when the two signals are dichotically combined over headphones, implicating a central mechanism that receives the inputs from the two ears, computes their invariance and differences, and translates these into a tonal percept. Therefore, unlike the Woods et al test which can be passed when listening to a single channel, to pass the HP test, participants must detect a target that is only perceived when L and R channels are fed separately to each ear (it is not present in each channel alone). Due to acoustic mixing effects, the percept is weak or absent when the stimuli are presented over loudspeakers. As a result, it should provide a more reliable screening tool. Similarly to Woods et al. (2017) we created a three-alternative forced-choice (3AFC) paradigm. Two intervals contain diotic white noise and the third (“target”) contains the dichotic stimulus that evokes the HP percept. Participants are required to indicate the interval that contains the hidden tone. This paradigm has the added attraction of being based on a detection task and so may, therefore, impose a lower load on working memory or other cognitive resources than the discrimination task of Woods et al. (2017).

To determine which test is more sensitive to headphone versus loudspeaker use, we directly compared the two approaches: The *Anti-Phase* (AP) test of Woods et al. (2017) and our new paradigm based on *Huggins Pitch* (HP). In Experiment 1 we used the Gorilla online platform (Anwyl-Irvine et al., 2020) to obtain performance from 100 “Trusted” participants (colleagues from across the auditory research community), who completed both tests over both loudspeakers and headphones. Importantly, each participant used their own computer setup and audio equipment, resulting in variability that is analogous to that expected for experiments conducted online (links to demos of each test are available in Materials and methods). In Experiment 2 we further tested the AP and HP screens using anonymous online participants. Participants in this group claimed to only use headphones to complete each test and we evaluated their performance using the profile of results we would expect for headphone and speaker use, based on Experiment 1.

Our results reveal that the HP test has a better diagnostic ability than the AP test to classify between headphones and loudspeakers. We also show that the classification performance can be improved further by combining the HP test with either the AP test or an alternative test based on beat perception (BT; Experiment 3).

## Experiment 1 – “Trusted” participant group

### Methods

#### Participants

114 “Trusted” participants were tested. Fourteen of these did not complete the full experiment (exited early mostly due to hardware issues e.g. incompatible loudspeakers or headphones). We report the results of the remaining 100 participants. Recruitment was conducted via email, inviting anyone who was over 18 and without known hearing problems to participate. The email was distributed to people we believed could be trusted to switch between headphones and loudspeakers when instructed to do so (e.g. via direct emails and via mailing lists of colleagues in the auditory scientific community). Participants were only informed of the general nature of the test which was to “assess remote participants’ listening environments”, with no reference to specific stimulus manipulations. Individual ages were not collected to help protect the anonymity of the participants. Grouped ages are presented in Table 1. Experimental procedures were approved by the research ethics committee of University College London [Project ID Number: 14837/001] and informed consent was obtained from each participant.

**Table 1.** Self-reported Participant Age Range in Experiment 1 (“Trusted” group), Experiment 2 (“Unknown” group, see below) and Experiment 3 (“Trusted” group).

|  | Experiment 1<br>(N =100) | Experiment 2<br>(N = 100) | Experiment 3<br>(N = 42) |
| --- | --- | --- | --- |
| **Age Bracket (% cases)** |  |  |  |
| 18-25 | 24 | 72 | 12 |
| 26-35 | 36 | 22 | 38 |
| 36-45 | 22 | 4 | 24 |
| 46-55 | 13 | 2 | 19 |
| 56-65 | 3 | - | 7 |
| Over 65 | 2 | - | - |

Data from this cohort may not represent ‘ground truth’ to the same extent as lab-based testing, but these participants were trusted to correctly implement the headphone and loudspeaker manipulation and their AP results were highly similar to the lab-based data collected by Woods et al. (2017), suggesting that data from this cohort was reliable.

#### Stimuli and Procedure

Gorilla Experiment Builder (www.gorilla.sc) was used to create and host the experiment online (Anwyl-Irvine et al., 2020). Participants were informed prior to starting the test that they would need access to both loudspeakers (external or internal to their computer) and headphones. The main test consisted of four blocks. Two blocks were based on the HP test and two on the AP test. Both HP and AP used a 3AFC paradigm (Figure 2). At the start of each block, participants were told whether to use loudspeakers or to wear headphones for that block. The blocks were presented in a random order using a Latin square design. In total the study (including instructions) lasted about 10 minutes, with each block (HP_headphone, HP_loudspeaker, AP_headphone, AP_loudspeaker) taking 1.5–2.5 minutes.

**Figure 2.**
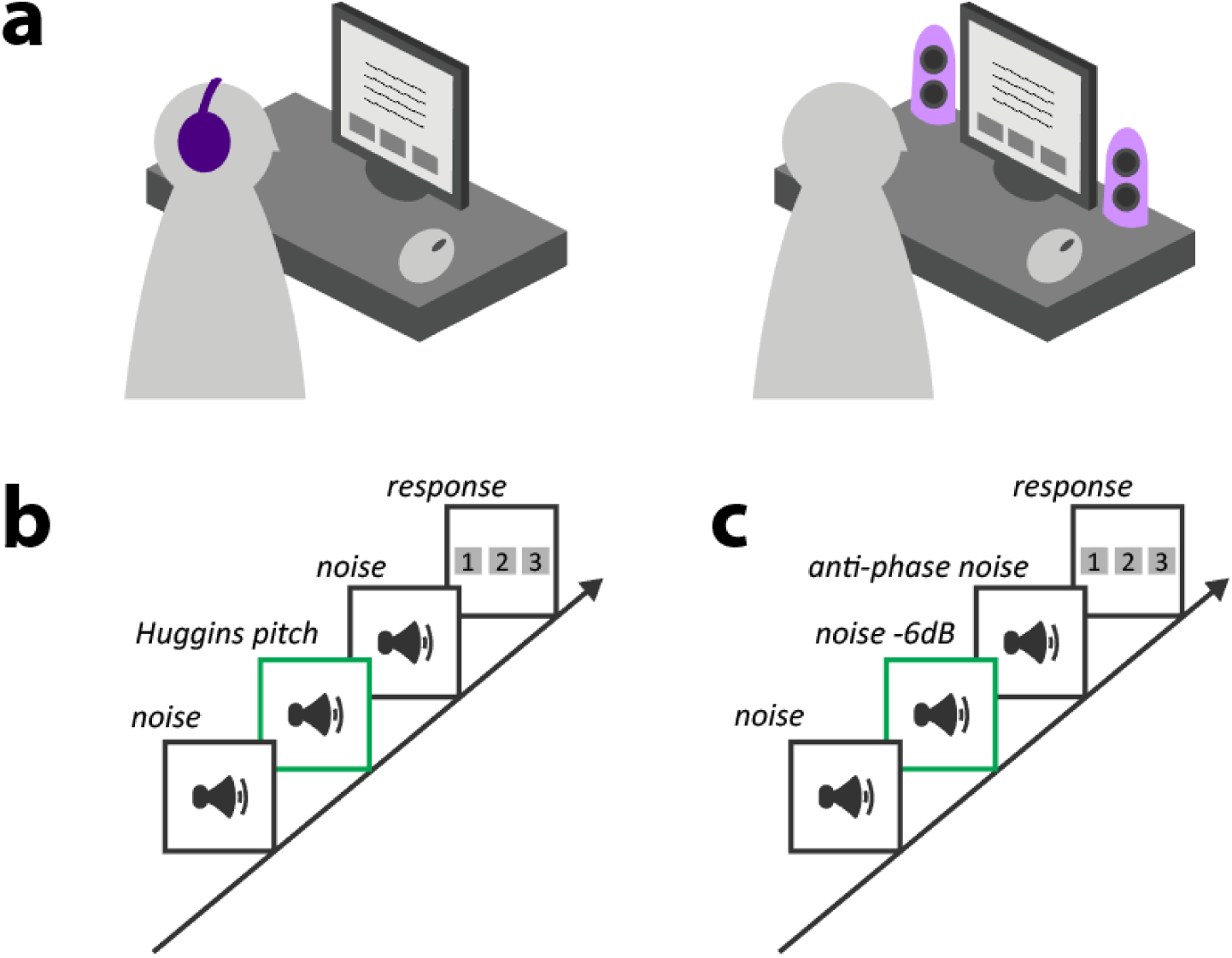
Schematic of the Test Design. (a) At the beginning of each block, participants were informed whether the upcoming test was to be performed whilst wearing headphones or via loudspeakers. Participants responded using a graphic user interface and computer mouse. The experiment was organised into four testing blocks: Huggins pitch (HP) test over headphones, HP test over loudspeakers, Anti-phase (AP) test over headphones, and AP test over loudspeakers. Test order was randomised across participants using a Latin square design. Both HP (b) and AP (c) tests used a 3AFC paradigm. For both tests, the example shows the target in the second position.

##### Volume Calibration

Every block began with a volume calibration to make sure that stimuli were presented at an appropriate level. For HP blocks a white noise was used; for AP blocks a 200-Hz tone was used. Participants were instructed to adjust the volume to as high a level as possible without it being uncomfortable.

##### HP Screening

The HP stimuli consisted of three intervals of white noise, each 1000 ms long. Two of the intervals contained diotically presented white noise (Figure 2). The third interval contained the HP stimulus. A center frequency of 600 Hz was used (roughly in the middle of the frequency region where HP is salient). The white noise was created by generating a random time sequence of Gaussian distributed numbers with a zero mean (sampling frequency 44.1 kHz, bandwidth 22.05 kHz). The HP signals were generated by transforming the white noise into the frequency domain and introducing a constant phase shift of 180° in a frequency band (± 6%) surrounding 600 Hz within the noise sample, leaving the amplitudes unchanged, and then converting the stimulus back to the time domain. The phase-shifted version was presented to the right ear, while the original version was delivered to the left ear (Yost et al., 1987). Overall, 12 trials were pre-generated offline (each with different noise segments; the position of the target uniformly distributed). For each participant, in each block (HP loudspeaker / HP headphones) 6 trials were randomly drawn from the pool without replacement.

The participant was told that they will “hear three white noise sounds with silent gaps inbetween. One of the noises has a faint tone within.” They were then asked to decide which of the three noises contained the tone by clicking on the appropriate button (1, 2, or 3).

##### AP Screening

The AP stimuli were the same as in Woods et al. (2017). They consisted of three 200-Hz tones (1000 ms duration, including 100 ms raised-cosine onset and offset ramps). Two of the tones were presented diotically: 1) the “standard”, and 2) the “target” which was the same tone at −6 dB relative to the standard. The third tone (the “foil”) had the same amplitude as the standard but was presented such that the left and right signals were in anti-phase (180°) (Figure 2). Listeners were instructed that “Three tones in succession will be played, please select the tone (1, 2, or 3) that you thought was the quietest”. As in the HP screening, for each participant, in each block (AP loudspeaker / AP headphones) 6 trials were randomly drawn from a pre-generated set of 12 trials.

Each screening test began with an example to familiarize the participants with the sound. The target in the example did not rely on dichotic processing but was simulated to sound the same as the target regardless of delivery device (for HP this was a pure tone embedded in noise; for AP two equal amplitude standards and a softer target were presented). Failure to hear the target in the example resulted in the participant being excluded from the experiment. Following the example, each block consisted of 6 trials. No feedback was provided, and each trial began automatically.

Test implementations are available in JavaScript https://sijiazhao.github.io/headphonecheck/ and via the Gorilla experimental platform https://gorilla.sc/openmaterials/100917. Stimuli can also be downloaded https://github.com/ChaitLabUCL/Headphones-test for implementation on other platforms. The AP code implementation from Woods et al. (2017) can be accessed via http://mcdermottlab.mit.edu/downloads.html. Versions of all tests can be previewed in a web browser using these URLs:

HP: https://sijiazhao.github.io/headphonecheck/headphonesTestHugginsPitch.html
AP: https://sijiazhao.github.io/headphonecheck/headphonesTestAntiPhase.html
BT: https://sijiazhao.github.io/headphonecheck/headphonesTestBeat.html

#### Statistical Analysis

We used signal detection theory to ask how well the two test types (HP and AP) distinguished whether participants were using headphones or loudspeakers. Accepting a user (i.e. deciding that they passed the test) at a given threshold (minimum number of correct trials) when they were using headphones was considered a “hit”, while passing that user at the same threshold when they were using loudspeakers was considered a “false alarm”. We used these quantities to derive a receiver operating characteristic (ROC; Swets, 1986) for each test type, enabling a comparison in terms of their ability to distinguish headphone versus loudspeaker use. As well as calculating the area under the ROC curve (AUC) as an overall sensitivity measure, we also report the sensitivity (*d*’) of the HP and AP tests at each of the thresholds separately. Note that “hits”, “false alarms”, and “sensitivity” here are properties of our tests (HP and AP) to detect equipment, not of the subjects taking those tests.

On the basis that a subject’s performance above chance should be a minimum requirement for them to be accepted under any selection strategy, we considered only thresholds (number of correct responses required to pass) of 3, 4, 5, and 6 trials out of 6. This approach also side-stepped the issue that the AP test over loudspeakers can result in below-chance performance, as evident in Figure 3 (light blue line does not show a chance distribution).

**Figure 3.**
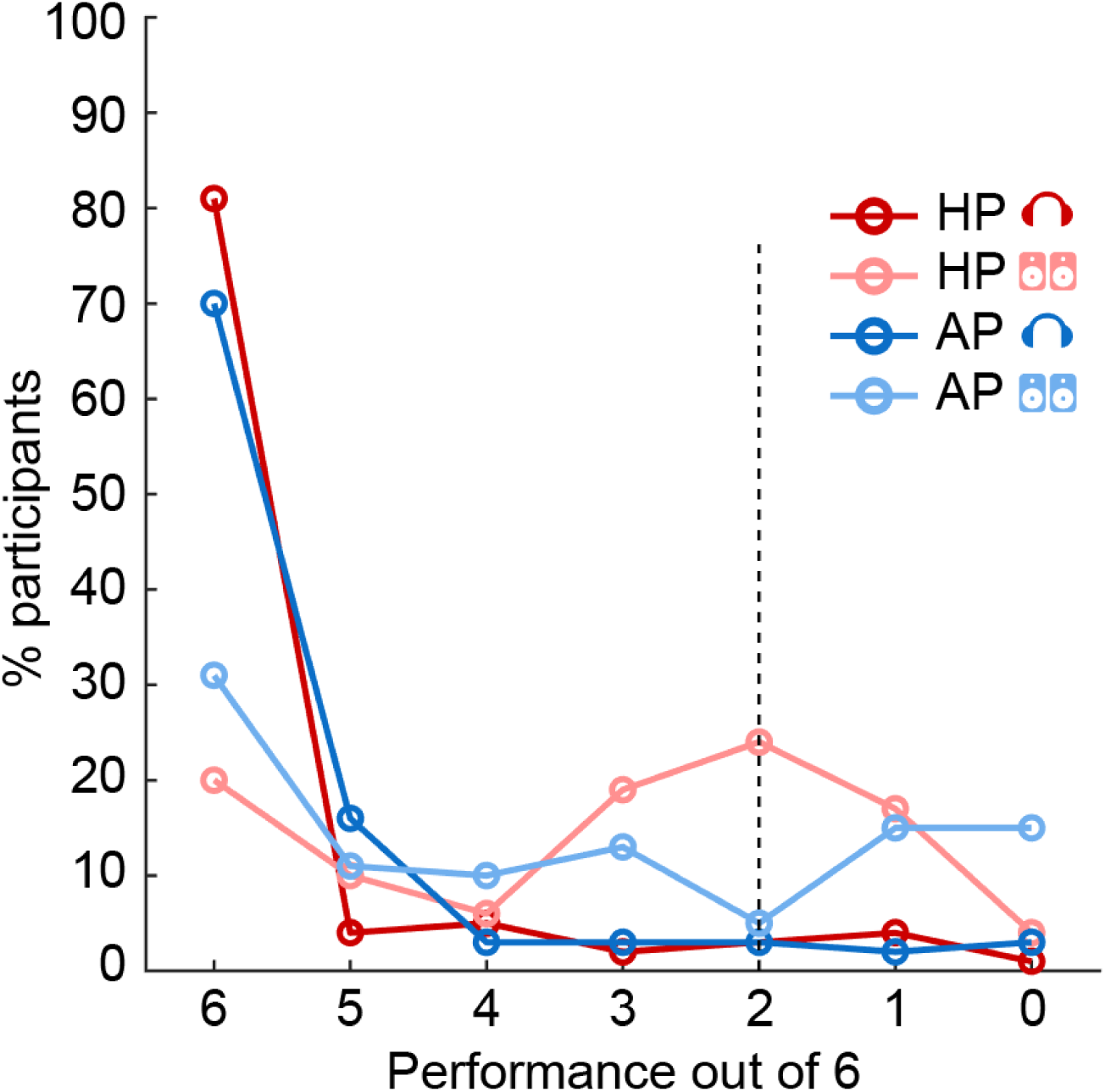
Distribution of Performance for Each Test Condition in the “Trusted” group (Experiment 1; N=100). The graph shows the proportion of participants (ordinate) at each level of performance (abscissa), ranging from a perfect score of 6/6 to 0/6 trials. The dashed black line indicates chance performance.

We additionally considered whether a combined test that made use of responses both to HP and AP trials would be more sensitive than either condition alone. Under this “Both” approach, subjects passed only if they met the threshold both for HP and AP trials.

We assessed statistical significance of differences in sensitivity (AUC) in two ways. First, we determined reliability of the results through bootstrapped resampling over subjects. For each of 1,000,000 resamplings we randomly selected 100 subjects with replacement from the pool of 100 subjects (balanced) and obtained a distribution of differences in the AUC for HP versus AP tests. We then determined the proportion of resamples for which the difference exceeded zero (separately for each direction of difference, i.e. HP minus AP, then AP minus HP), and accepted the result as significant if this was greater than 97.5% in either direction (two tailed; *p* < 0.05). The other method we used to assess statistical significance of differences of interest was with respect to a null distribution obtained through relabelling and permutation testing. For each of 1,000,000 permutations we randomly relabelled the two headphone condition scores for each of the 100 subjects as HP or AP, and similarly for the two loudspeaker scores. We then calculated the AUC at each threshold for these permuted values. This generated a null distribution of AUC differences that would be expected by chance. We then determined the proportion of scores in these null distributions that exceeded the observed difference in either direction and accepted the result as significant if this was less than 2.5% in either direction (two tailed; *p* < 0.05). Identical procedures were used to test for differences between the “Both” approach and each of the HP and AP methods.

### Results

#### Distribution of performance for each screening test

Figure 3 presents a distribution of performance across participants and test conditions. The x axis shows performance (ranging from a perfect score of 6/6 to 0/6). Chance performance (dashed black line) is at 2. The performance on the AP test with headphones (dark blue line) generally mirrored that reported in Woods et al. (2017), except that the pass rate in the headphones condition (70%) is substantially lower than in their controlled lab setting data (100%). This is likely due to the fact that the “Trusted” participants in the present experiment completed the test online, thereby introducing variability associated with specific computer/auditory equipment. Performance on the AP test with loudspeakers (light blue line) was also similar to that expected based on the results of Woods et al. (2017). Some participants succeeded in the test over loudspeakers (30% at 6/6). Notably, and similarly to what was observed in Woods et al. (2017), the plot does not exhibit a peak near 2, as would be expected by chance performance in a 3AFC task, but instead a *trough,* consistent with participants mistaking the phase shifted “foil” for the “target”. For the HP test, a chance distribution is clearly observed in the loudspeaker data (peak at 2, light red line). There is an additional peak at 6, suggesting that some participants (20% at 6/6) can detect Huggins Pitch over loudspeakers. In contrast, performance using headphones for HP (dark red line) shows an “all-or-nothing” pattern with low numbers for performance levels below 6/6, consistent with HP being a robust percept over headphones (Akeroyd et al., 2001).

#### Ability of each screening test to distinguish between headphone vs. loudspeaker use

We derived the receiver operating characteristic (ROC) for each test, plotting the percentage of participants who passed at each above-chance threshold while using headphones (“hits”, y-axis) or loudspeakers (“false alarms”, x-axis) (Figure 4a). The area under the curve (AUC) provides a measure of how well each test type distinguishes between headphone versus loudspeaker use. The AUC for HP (.821) was significantly larger than that for AP (.736) (bootstrap resampling: *p* =. 022, permutation test: *p*=. 018). This suggested that the HP test overall provides better overall sensitivity (i.e., maximizing the headphones pass rate, while reducing the proportion of listeners who pass using loudspeakers). This is also illustrated in Figure 4b which plots *d*’ at each threshold. The maximum *d*’ reached is ∼1.7, consistent with medium sensitivity at the highest threshold (6/6). At this threshold HP will correctly detect 81% of the true headphone users, but also pass 20% of loudspeaker users, whereas AP will detect 70% of the headphone users, but also pass 31% of loudspeaker users; for threshold of 5/6 the values are 85%/30% for HP and 86%/42% for AP.

**Figure 4.**
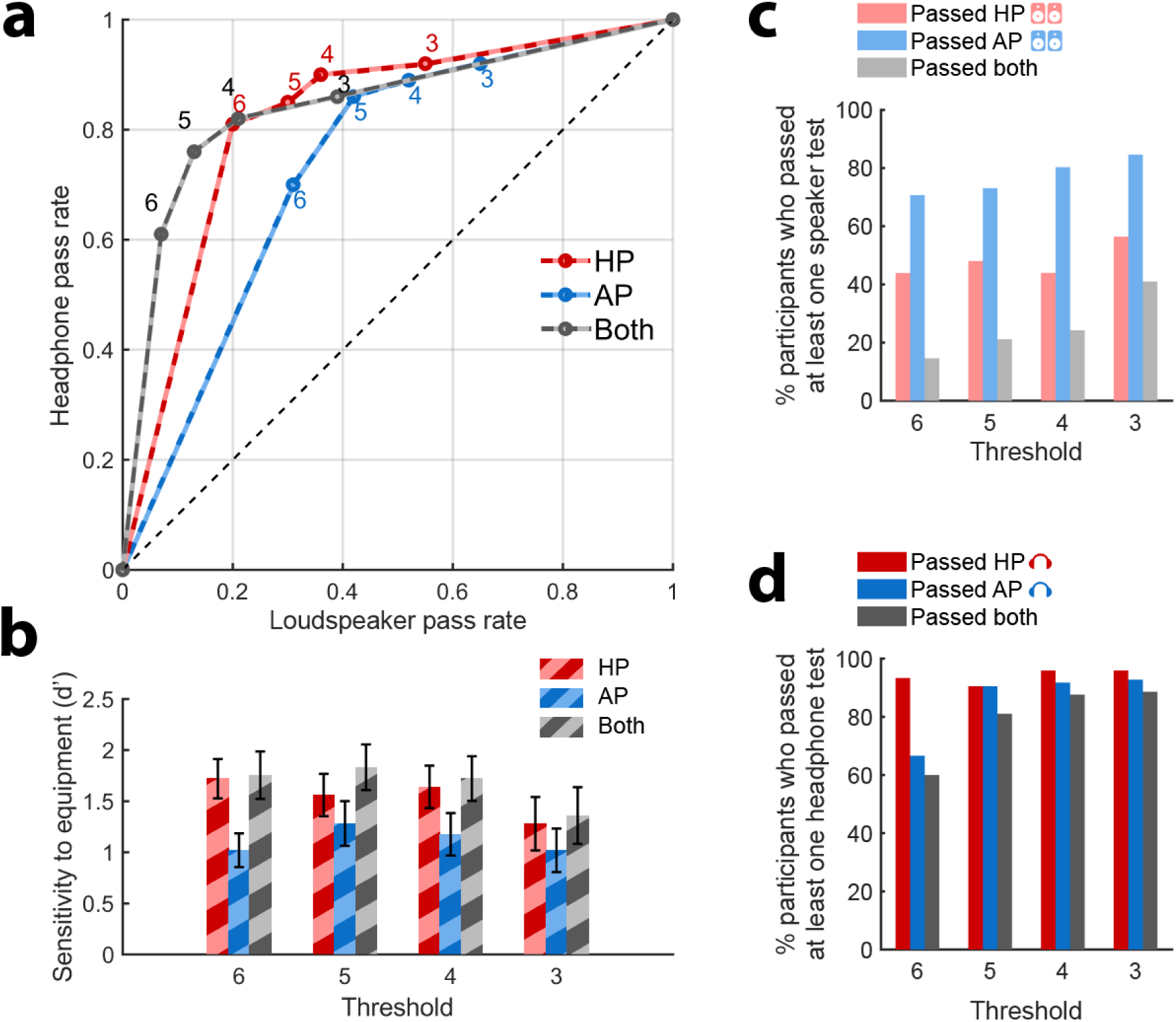
Ability of HP, AP and a Combined Test (“Both”) to Distinguish Between Headphone and Loudspeaker Users (N = 100). (a) ROC curves. The proportion of participants passing at each above-chance threshold (3,4,5,6 /6 labelled next to each data point) while using headphones (“hits”, y-axis) or loudspeakers (“false alarms”, x-axis) for HP, AP or a combined test (“Both”). (b) Sensitivity (d’) at each threshold. Error bar = 1 std bootstrap with 10,000 iterations. (c) Pass rates with loudspeakers at each threshold, plotted relative to the total number of participants who passed at least one of the loudspeaker tests. (d) Pass rates with headphones. Whilst a large proportion of participants pass both AP and HP tests over headphones (dark grey bars in d), only a small proportion pass both tests over loudspeakers (light grey bars in c).

We also plotted the ROC and sensitivity for a “Both” approach that required participants to reach the threshold both for the HP and AP tests. The AUC for Both was. 844 and significantly higher than for AP (bootstrap resampling *p* <. 001, permutation test: *p* =. 014) but not for HP (bootstrap resampling: *p =*. 279, permutation test: *p* =. 979). Given the additional time that would be required compared to running HP alone, the lack of significant difference over HP suggests that the combined test is not generally a worthwhile screening approach. However, if the experiment is such that headphone use is critical then using the combined test will reduce the loudspeaker pass rate from 20% to 7% but at the expense of rejecting 40% of headphone users. This is illustrated in Figure 4c, which plots the proportion of listeners who pass the AP and HP tests over loudspeakers (relative to the number of subjects who pass at least one test over loudspeakers). For each threshold, the proportion of listeners who pass the AP test over loudspeakers is larger than that for HP (Figure 4c). The proportion of listeners who pass both loudspeaker tests is very low, consistent with the fact that the conditions that promote passing the HP test over loudspeakers (listeners close to and exactly between the loudspeakers such that left and right ears receive primarily the left and right channels, respectively) are antithetical to those that yield better AP performance. Therefore, combining the two tests will substantially reduce the number of people who pass using loudspeakers. In contrast to the performance with loudspeakers, most participants passed both the HP and AP tests when using headphones (Figure 4d). The higher HP pass rates in Figure 4d may stem from the fact that the audio equipment used by a large proportion of participants have some bleed between L and R channels such that the HP test is still passable but performance on the AP test is affected more severely. Therefore, combining both tests (‘BOTH’) can provide a strict test of stereo headphone use. We return to this point in Experiment 3, below.

## Experiment 2 - “Unknown” online group

We probed performance on the AP and HP tests in a typical online population. This time, participants were unknown to us, recruited anonymously and paid for their time. We informed participants that headphones had to be worn for this study and sought to determine whether the pass rate would be similar to that in the “Trusted” cohort.

### Methods

#### Participants

We recruited online participants via the Prolific recruitment platform (prolific.co). Of the 103 participants who were tested, three were unable to hear one of the example sounds and left the study early, leaving a total of 100 participants. Participants were paid to complete the 5-7-minute study. We specified that they should not accept the study if they had any known hearing problems. No additional exclusion criteria were applied to this sample in order to obtain a broad range of participants. Reported ages are provided in Table 1 (middle panel). Experimental procedures were approved by the research ethics committee of University College London [Project ID Number: 14837/001] and informed consent was obtained from each participant.

#### Stimuli and procedure

Stimuli and procedure were the same as in Experiment 1 except participants were instructed to only use headphones, thus completing each screening test, HP and AP, once. The instructions stressed that headphones must be worn for this experiment.

## Results

Figure 5 plots the performance (black lines) observed for the “Unknown” online group. Overall, the performance patterns were different from the performance using headphones obtained from the “Trusted” group, suggesting that a proportion of listeners may not have heeded the instructions to use headphones, or used low quality equipment. In particular, there was a ∼10% greater number of participants getting 6/6 with the AP vs. HP test, which is the reverse of what was seen in the “Trusted” group with headphones. This adds support to the results of Experiment 1 that suggest there is a higher false positive rate with the AP test.

**Figure 5.**
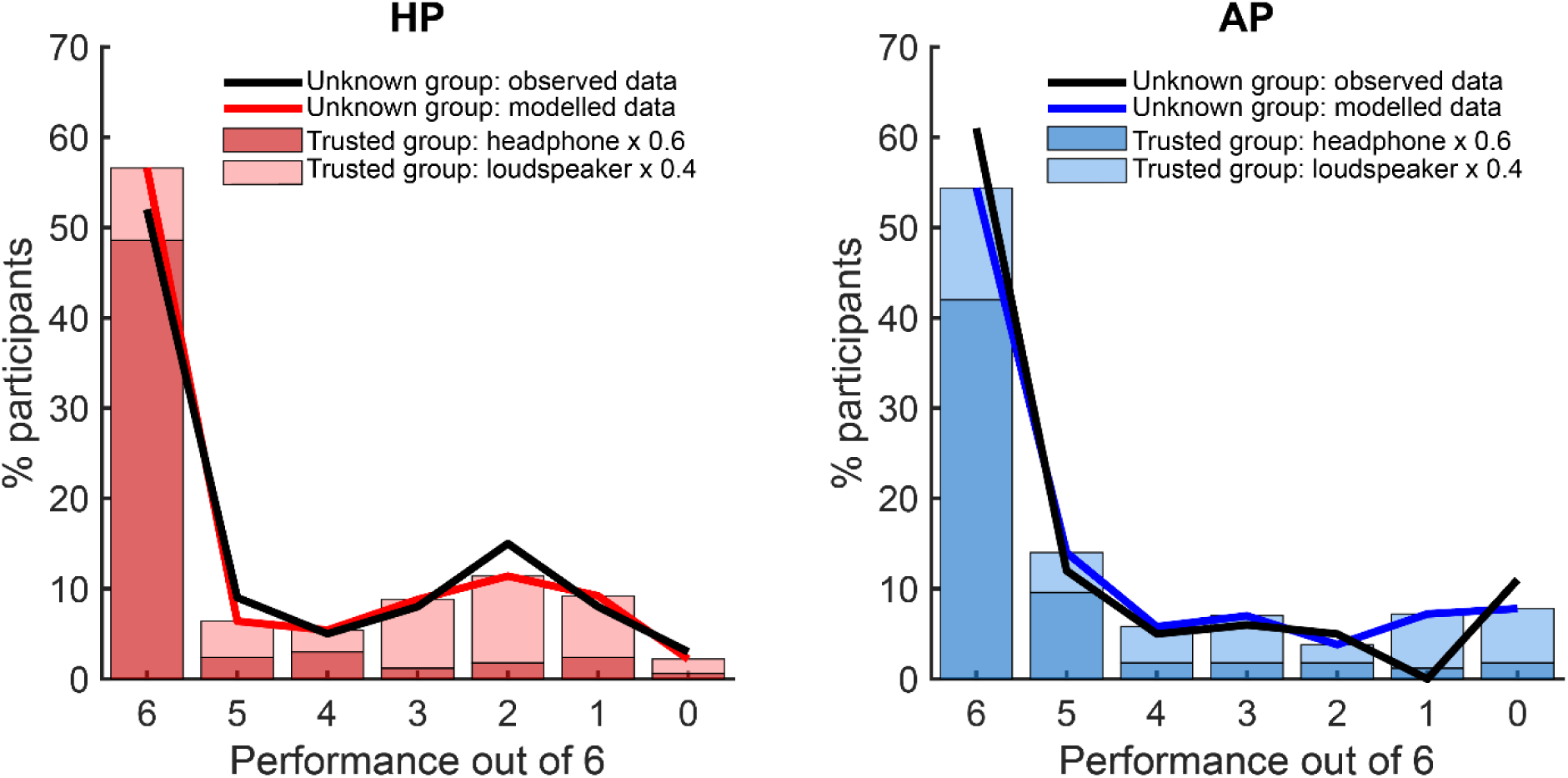
Performance in the “Unknown” Group (Experiment 2; N=100). The black solid line illustrates the performance observed in the “Unknown” group in the HP (left) and AP (right) tests. The red/blue lines and stacked bars illustrate the result of modelling to determine the likely proportion of subjects in the “Unknown” group who actually used headphones. The stacked bars indicate the product of the proportion of participants in the “Trusted” group at each performance level with the relevant coefficient from the best fitting model (0.6 for headphones and 0.4 for loudspeakers, fixed across HP and AP). The distribution of observed performance in the “Unknown” group matched the modelled data well for both HP and AP tests. This indicates that only roughly 60% of participants in the “Unknown” group showed performance that was consistent with headphone use.

To estimate the proportion of online participants that actually used headphones, we assumed that the distribution of online scores for each test type could be explained as a linear combination of the distributions of headphone and loudspeaker scores from the same test type in the “Trusted” group (Experiment 1). We used a simple model with a single parameter, *propH*, for the proportion of headphone users. For values of *propH* varying from 0 to 1 in increments of. 01 we multiplied the distribution of Experiment 1 HP headphone scores by *propH*, and summed these two values, giving a modelled distribution of Experiment 2 HP scores for each value of *propH*. We repeated the same process for AP scores. We then compared the modelled and observed distributions and selected the value of *propH* that minimised the sum of squared errors across both HP and AP scores. This analysis yielded an estimate that 40% of users in Experiment 2 likely did not use headphones (or had unsuitable equipment/completed the test in a noisy setting), demonstrating the importance of running an objective screen.

## Experiment 3 – Combination testing: Huggins Pitch and Beat Test

In experiment 1 we demonstrated that combining HP and AP tests can provide greater selectivity for headphones than using the HP test alone. In experiment 3, we examined the use of a different test based on beat stimuli (BT), that can potentially be combined with the HP test to provide better selectivity for headphone user. Monaural beats are perceived when two tones of similar but nonidentical frequencies (e.g. 1800 Hz and 1830 Hz) are presented simultaneously. The listener perceives fluctuations or beats produced by amplitude modulation whose rate is equal to the difference of the two frequencies (Oster, 1973). A related binaural phenomenon occurs when the tones are presented to each ear separately. This “binauaral beat” is perceived due to central interference between the two frequencies. However, due to the phase locking limits on binaural processing, binaural beats are only perceived for frequencies lower than 1000-1500 Hz (Licklider et al., 1950; Perrott & Nelson, 1969; Rutschmann & Rubinstein, 1965). In addition, binaural beats are only salient for relatively small frequency differences (< 10 Hz). We take advantage of this difference between diotic and dichotic stimulation to create a test of L and R channel independence.

The stimulus (Figure 6) consists of simultaneous presentation of two pure tones (“pair”) of frequencies f1 and f2. f1 is randomly drawn from between 1800 and 2500 Hz and f2 is set to f1+30 Hz. In each trial the listener is presented with three intervals, each containing a pair of pure tones, and must determine which interval was the smoothest. Two of the pairs are presented diotically (“standards”; Figure 6a) and should be associated with a strong perception of a beat at 30 Hz. In the other pair (“target”; Figure 6b), the tones are presented dichotically, one to each ear. Because the frequencies are above the phase locking limit, and because the frequency difference is higher than the typical limit of binaural beats, the stimulus should not lead to a binaural beat percept and will therefore be heard as “smooth” over headphones. However, over loudspeakers the left and right signals interact before reaching the listener’s ears to create a monaural beat percept, making the target indistinguishable from the standards.

**Figure 6.**
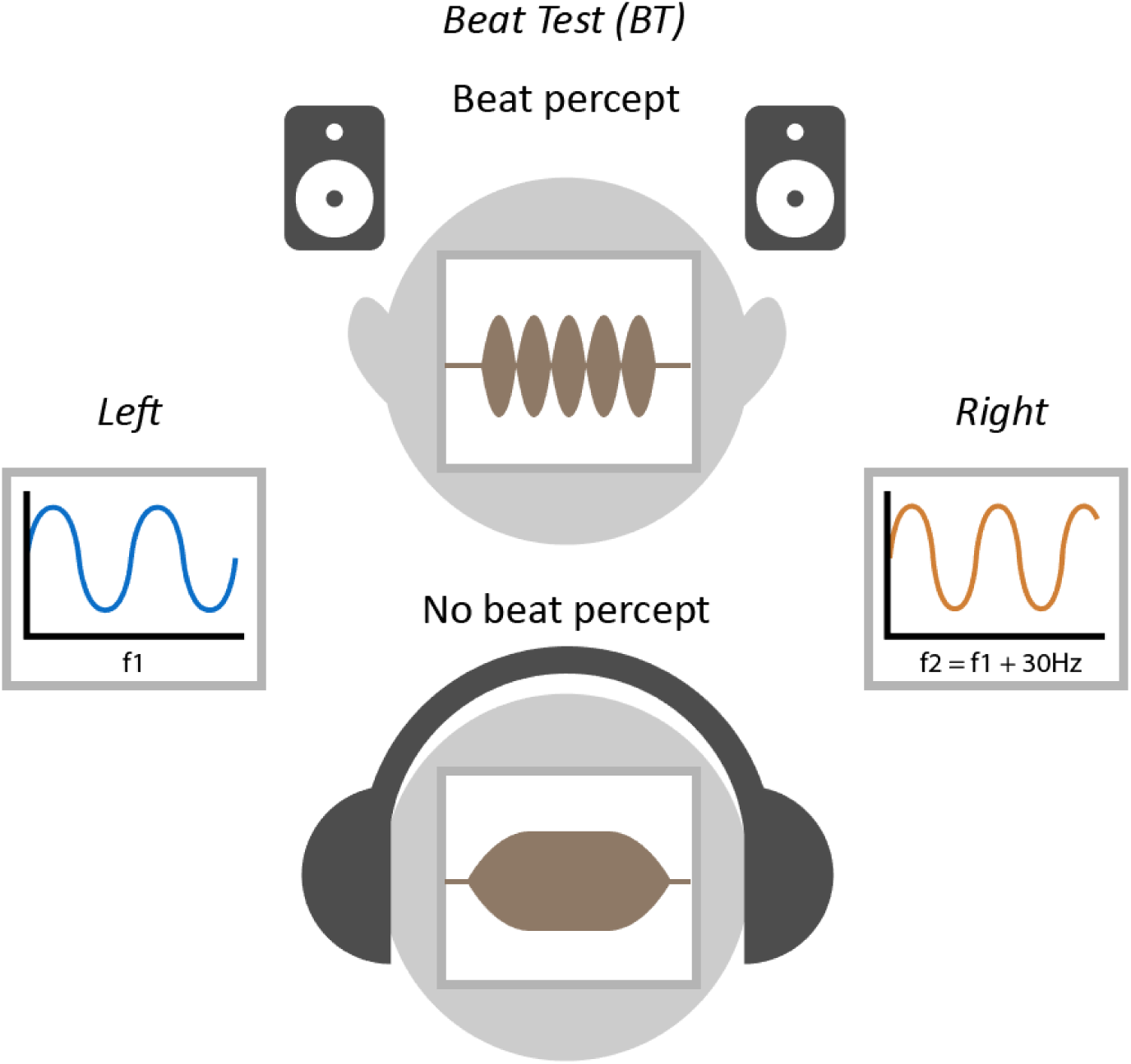
Schematic of stimuli used for the BT test. When f1 (f1>1500 Hz) and f2 (f2=f1+ 30 Hz) are presented dichotically over headphones, a smooth tone is heard (bottom). When f1 and f2 are presented diotically through headphones or through channel interference via loudspeakers (top), then a monaural beat percept is heard. In the BT test the percept without the beat is the target which is foiled when heard over loudspeakers.

This test is similar to the AP test in that it relies on channel interference over loudspeakers and therefore suffers from analogous constraints, including being affected by the specific positions of the loudspeakers relative to the listener. Furthermore, similarly to AP, it is possible to pass the BT test when listening over a single channel (e.g. when listening through a single ear bud), so that it cannot be used as a headphone screening test on its own. However, the BT test has several advantages over the AP test which might make it a more efficient complement for HP. Notably the target (“smooth tone”) and distractors (“beats”) are more perceptually distinct than the level difference used in the AP test, which might result in better discrimination performance. Furthermore, the robust binaural effect is expected to lead to lower pass rates over loudspeakers. We therefore reasoned that the BT test, when used in combination with the HP test, may provide a sensitive probe of headphone use than that demonstrated in Experiment 1 (HP+AP).

We therefore recruited a further group of “Trusted” participants who completed the HP and BT tests over loudspeakers and headphones.

### Participants

42 “Trusted” participants were tested. Recruitment was conducted in the same way described for Experiment 1. Grouped ages are presented in Table 1. Experimental procedures were approved by the research ethics committee of University College London [Project ID Number: 14837/001] and informed consent was obtained from each participant.

### Stimuli and procedure

The paradigm was identical to the 3-AFC test used in experiment 1 except that participants completed the beat (BT) test in place of the AP test. The BT stimuli consisted of three intervals, each 1000 ms long. Two of the intervals contained diotically presented tone pairs. The frequency of the first tone (f1) was randomly drawn from 1800-2500Hz. The frequency of the second tone (f2) was set to f1+30 Hz. The third interval contained a dichotically presented tone pair. All tones were gated with a 5-ms raised-cosine onset and offset ramps. To reduce reliance on any loudness cues the amplitude of each interval was randomly roved to result in relative differences of 0-4 dB. 12 trials were pre-generated offline (with the position of the target uniformly distributed). For each participant in each block, 6 trials were randomly drawn from the pool without replacement. In the BT test, listeners were informed that they would hear three sounds in succession and asked to detect the sound (1,2 or 3) that they thought was the “smoothest”. Statistical analysis was performed in the same way as for experiment 1.

### Results

ROC and AUC analysis was conducted in the same way as for experiment 1. The results are presented in Figure 7. Figure 7a plots the derived ROC curves for the HP test, BT test, and the combined test (‘both’; HP+BT). To compare HP performance across experiments 1 and 3 we used a bootstrap resampling procedure (Figure 7d) whereby subsets of 42 participants were repeatedly (N=1000) sampled (without replacement) from the experiment 1 data set (N=100). This analysis demonstrated that the ROC for HP obtained in experiment 3 is in line with that observed in experiment 1. The resampling data clearly shows that there is substantial variability across participants, which would be expected given that online users will have different sound delivery setups and operate in different environments.

**Figure 7.**
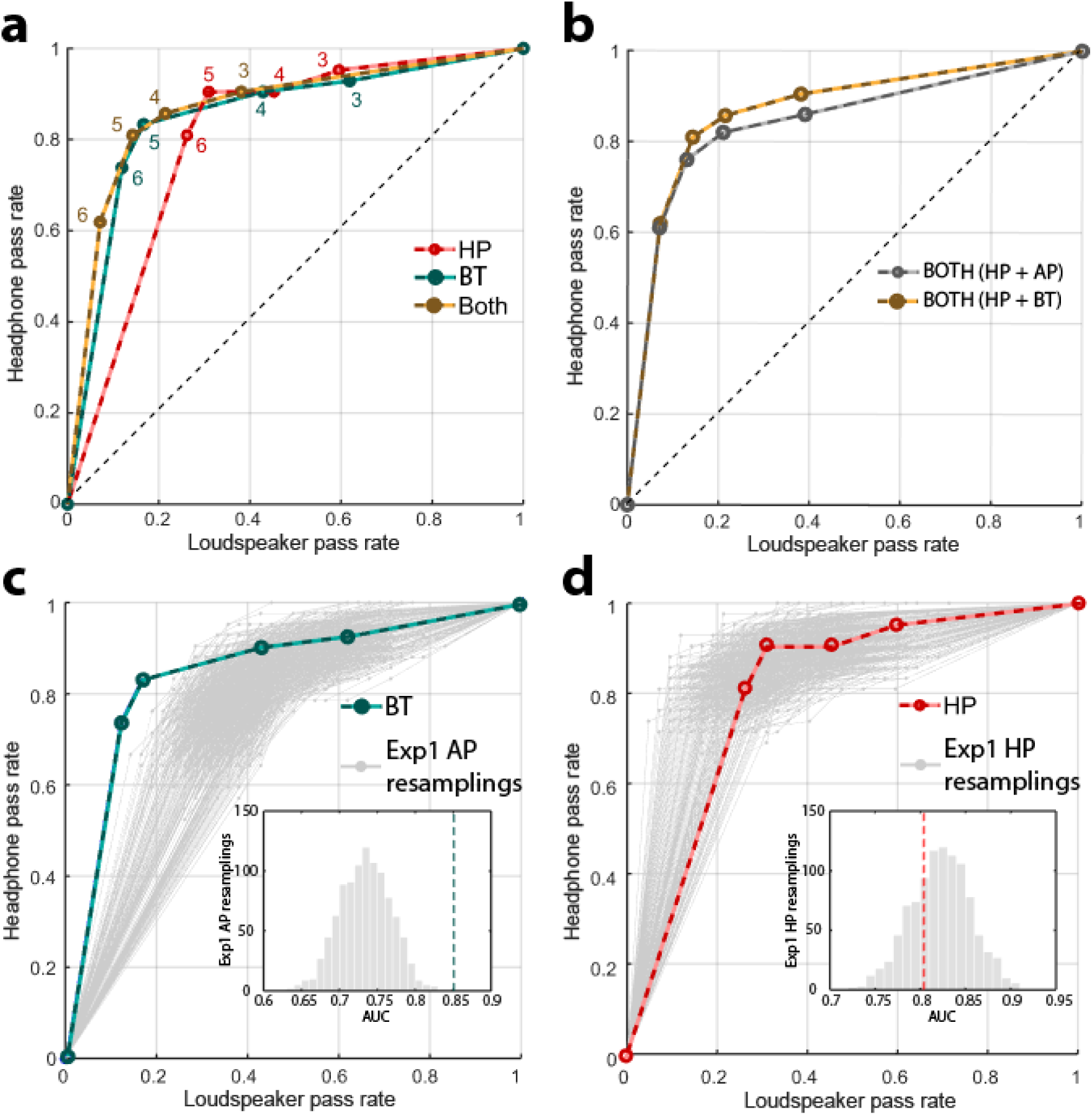
Sensitivity of HP and BT from experiment 3 (N = 42) and comparison with experiment 1. a) ROC curves. The proportion of participants passing at each above-chance threshold (3,4,5,6 /6 labelled next to each data point) while using headphones (“hits”, y-axis) or loudspeakers (“false alarms”, x-axis) for HP, BT or a combined test (“Both”). B) Comparison of combined tests of HP with either AP or BT. C) Comparison of BT and AP: The green line shows the ROC curve for the BT test. Grey lines show 1000 resamplings of 42 subject datasets from the AP test in experiment 1. s. D) Comparison of HP from experiment 1 and 3: The red line shows the ROC curve for the HP test from experiment 3, the grey lines show 1000 resamplings of 42 subject datasets from the HP test in experiment 1.

A similar bootstrap analysis was used to compare BT performance in experiment 3 to AP performance in experiment 1 (Figure 7c). The results reveal that, as hypothesized, the BT test was better able to distinguish between headphone vs. loudspeaker users than the AP test (Figure 7C).

AUC analysis indicated that the AUC for the combination of HP and BT was larger than that for HP (Bootstrap: *p*=.028), confirming that the combined test has better diagnostic ability than HP alone. However, notably, the ROC curves for the combination of HP+BT vs. that of HP+AP were very similar, with a marked overlap at higher thresholds (Figure 7b), suggesting it is difficult to improve on the previously demonstrated performance of a ∼7% false positive rate and ∼60% hit rate. As stated above, the relatively low true positive rate, even among our “trusted” participant group, may reflect the fact that a large proportion of the population is using audio equipment (e.g. sound card or headphones) that has some crosstalk between L and R channels, resulting in participants failing the tests despite using headphones.

### Discussion

We sought to develop an efficient headphone screening test for use with online auditory experiments that is easy to explain to listeners, quick to administer (< 3 mins) and which has a high selectivity for headphone users. We devised a new test (HP) based on a perceptual phenomenon that can only be detected when stimuli are presented dichotically. This detection test was contrasted with an existing test (AP). The analyses we reported demonstrate that HP has higher selectivity for headphone users than AP, rendering it a compelling alternative to the existing screening method. That it is based on a detection rather than a discrimination task, and therefore less dependent on working memory, further adds to its appeal.

We note that all our estimates are based on the “Trusted” participant group. However, this cohort (primarily from a network of colleagues and our scientific community) may not be fully representative of the general online participant population. For instance, it is conceivable that they possess higher quality equipment or were more motivated than the average online participant. In general, it is prudent to treat the “Trusted” group data as reflecting the best-case scenario with actual performance probably somewhat lower in the general population. Importantly, the test is designed to distinguish between participants who are using stereo headphones (i.e. where the left and right channels are independently delivered to the left and right ear, respectively) from those listening without headphones (where typically the left and right channels will interact in some way before reaching the listeners’ ears). Though the screen is not designed to be sensitive to other aspects of the listener’s environment per se, headphone users may nonetheless fail the test if the quality of the equipment is very low, their environment is particularly noisy or they have a hearing impairment.

Overall, we conclude that the HP test is a powerful tool to screen for headphone use in online experiments. We have made our implementation openly available and ready for use via JavaScript and Gorilla (gorilla.sc). The test consists of 6 trials and, based on the ROC analysis, our recommendation is to use a threshold of 6/6. Lower thresholds will result in a similar *d’* but will pass a larger proportion of loudspeaker users.

The HP test passes only 80% of “true” headphone users and fails to reject 20% of loudspeaker users. Failing the test over headphones could be attributable to poor quality equipment (e.g. crosstalk between left and right channels), background noise, or hearing impairment. Conversely, those subjects who pass with loudspeakers might be optimally spatially positioned (e.g. equally between the two loudspeakers for HP). In situations where it is important to reach a high level of certainty that the participant is using headphones (e.g., where stimuli involve a dichotic presentation or a spatial manipulation) the HP test can be combined with the BT test (experiment 3). This will yield a false positive rate of ∼7%.

That the combined test rejects ∼40% of “true” headphone users is an important observation and suggests that many household sound delivery systems suffer from bleed between left and right channels (introduced either by the sound card or headphones). This will reduce performance on the HP task, and more extensively so on the BT task, which is less robust to crosstalk between channels. Whilst some crosstalk may be inconsequential for most experiments that employ diotic sound presentation, studies that rely on specifically controlled stereo stimulation may be severely affected. The combined HP+BT test is a useful filter for such situations.

Overall, the rapid tests we have validated here can effectively aid researchers in confirming the listening environment of their participants thereby reaping the benefits of using online experimental platforms whilst controlling (at least certain aspects of) data quality.

## Funding

This work was funded by BBSRC grant [BB/P003745/1] to MC, a Wellcome Trust grant [213686/Z/18/Z] to AEM and supported by NIHR UCLH BRC Deafness and Hearing Problems Theme. AJB is supported by Wellcome Trust grant [091681]. KCP is supported by an ERC grant [771550-SOUNDSCENE]. AJO is supported by NIH grants (R01 DC005216 and R01 DC012262).

## Acknowledgments

We are grateful to Anahita Mehta, Gordon Mills and Fred Dick for comments and advice and to our colleagues across the international auditory research community for participating in the online experiments and providing feedback.

## Competing Interests

The authors declare no competing interests.

## Open Practices Statements

Data, analysis code and stimuli are available at https://github.com/ChaitLabUCL/Headphones-test. Code implementing the three headphone screening tests in JavaScript can be downloaded from the project website [https://sijiazhao.github.io/headphonecheck/]. Gorilla (gorilla.sc) versions of the three screening tests are available through Gorilla Open Material https://gorilla.sc/openmaterials/100917. Neither experiment was pre-registered.

## Notes

https://gorilla.sc/openmaterials/100917

https://sijiazhao.github.io/headphonecheck/

### Competing Interest Statement

The authors have declared no competing interest.

### Summary of Updates

This version was revised based on reviewer comments and also demonstrates that combining the Huggins Pitch test with an additional test, based on binaural beats, can yield a stricter measure (fewer false positives) of headphone use.

## References

Akeroyd, M. A., Moore, B. C. J., & Moore, G. A. (2001). Melody recognition using three types of dichotic-pitch stimulus. The Journal of the Acoustical Society of America, 110(3), 1498–1504. https://doi.org/10.1121/1.1390336

Anwyl-Irvine, A. L., Massonnié, J., Flitton, A., Kirkham, N., & Evershed, J. K. (2020). Gorilla in our midst: An online behavioral experiment builder. Behavior Research Methods, 52, 388–407. https://doi.org/10.3758/s13428-019-01237-x

Assaneo, M. F., Ripollés, P., Orpella, J., Lin, W. M., de Diego-Balaguer, R., & Poeppel, D. (2019). Spontaneous synchronization to speech reveals neural mechanisms facilitating language learning. Nature Neuroscience, 22(4), 627–632. https://doi.org/10.1038/s41593-019-0353-z

Chait, M., Poeppel, D., & Simon, J. Z. (2006). Neural response correlates of detection of monaurally and binaurally created pitches in humans. Cerebral Cortex, 16(6), 835–858. https://doi.org/10.1093/cercor/bhj027

Clifford, S., & Jerit, J. (2014). Is There a Cost to Convenience? An Experimental Comparison of Data Quality in Laboratory and Online Studies. Journal of Experimental Political Science, 1(2), 120. https://doi.org/10.1017/xps.2014.5

Cramer, E. M., & Huggins, W. H. (1958). Creation of Pitch through Binaural Interaction. Journal of the Acoustical Society of America, 30(5), 412–417. https://doi.org/10.1121/1.1909628

Kell, A. J. E., Yamins, D. L. K., Shook, E. N., Norman-Haignere, S. v., & McDermott, J. H. (2018). A Task-Optimized Neural Network Replicates Human Auditory Behavior, Predicts Brain Responses, and Reveals a Cortical Processing Hierarchy. Neuron, 98(3), 630–644. https://doi.org/10.1016/j.neuron.2018.03.044

Lavan, N., Knight, S., Hazan, V., & McGettigan, C. (2019a). The effects of high variability training on voice identity learning. Cognition, 193, 104026. https://doi.org/10.1016/j.cognition.2019.104026

Lavan, N., Knight, S., & McGettigan, C. (2019b). Listeners form average-based representations of individual voice identities. Nature Communications, 10(1), 1–9. https://doi.org/10.1038/s41467-019-10295-w

Licklider, J. C. R., Webster, J. C., & Hedlun, J. M. (1950). On the Frequency Limits of Binaural Beats. Journal of the Acoustical Society of America, 22(4), 468–473. https://doi.org/10.1121/1.1906629

McPherson, M. J., & McDermott, J. H. (2018). Diversity in pitch perception revealed by task dependence. Nature Human Behaviour, 2(1), 52–66. https://doi.org/10.1038/s41562-017-0261-8

Oster, G. (1973). Auditory beats in the brain. *Scientific American*. https://doi.org/10.1038/scientificamerican1073-94

Perrott, D. R., & Nelson, M. A. (1969). Limits for the Detection of Binaural Beats. The Journal of the Acoustical Society of America, 46(6B), 1477–1481. https://doi.org/10.1121/1.1911890

Rodd, J. (2019). How to Maintain Data Quality When You Can’t See Your Participants. APS Observer, 32(3). https://www.psychologicalscience.org/observer/how-to-maintain-%09data-quality-when-you-cant-see-your-participants

Rutschmann, J., & Rubinstein, L. (1965). Binaural Beats and Binaural Amplitude-Modulated Tones: Successive Comparison of Loudness Fluctuations. Journal of the Acoustical Society of America, 38(5), 759–768. https://doi.org/10.1121/1.1909802

Slote, J., & Strand, J. F. (2016). Conducting spoken word recognition research online: Validation and a new timing method. Behavior Research Methods, 48(2), 533–566. https://doi.org/10.3758/s13428-015-0599-7

Swets, J. A. (1986). Indices of Discrimination or Diagnostic Accuracy. Their ROCs and Implied Models. Psychological Bulletin, 99(1), 100. https://doi.org/10.1037/0033-2909.99.1.100

Woods, A. T., Velasco, C., Levitan, C. A., Wan, X., & Spence, C. (2015). Conducting perception research over the internet: A tutorial review. PeerJ, 3, e1058. https://doi.org/10.7717/peerj.1058

Woods, K. J. P., & McDermott, J. H. (2018). Schema learning for the cocktail party problem. Proceedings of the National Academy of Sciences of the United States of America, 115(14), E3313–E3322. https://doi.org/10.1073/pnas.1801614115

Woods, K. J. P., Siegel, M. H., Traer, J., & McDermott, J. H. (2017). Headphone screening to facilitate web-based auditory experiments. *Attention*, Perception, and Psychophysics, 79(7), 2064–2072. https://doi.org/10.3758/s13414-017-1361-2

Yost, W. A., Harder, P. J., & Dye, R. H. (1987). Complex spectral patterns with interaural differences: dichotic pitch and the ‘central spectrum’. In: Auditory processing of complex sounds No Title. In *Auditory processing of complex sounds* (pp. 190–201).

Zhao, S., Yum, N. W., Benjamin, L., Benhamou, E., Yoneya, M., Furukawa, S., Dick, F., Slaney, M., & Chait, M. (2019). Rapid Ocular Responses Are Modulated by Bottom-up-Driven Auditory Salience. Journal of Neuroscience, 39(39), 7703–7714. https://doi.org/10.1523/JNEUROSCI.0776-19.2019

